# Interaction of Ricin and RCA-I lectin with Human Serum Glycoproteins and Lung Cells

**DOI:** 10.1101/2025.05.08.652972

**Authors:** J.A. Smith, O.A. Vanderpuye

## Abstract

The plant protein ricin is one of the most toxic substances known and has been used in crimes internationally. Ricin binds to cell surface galactose as on glycoproteins and becomes endocytosed to kill cells. Blood contains high levels of galactosylated proteins which could bind to ricin and the related protein RCA-I from the same plant Ricinus communis. While blood proteins would be among targets for injected ricin, inhaled ricin would also bind to lung epithelial cells. Ricin and RCA-I are reported to have very similar or identical binding specificity. Thus, it is useful to increase the knowledge of proteins in blood and A549 cells that bind to ricin and RCA-I.

Ricin and RCA-I staining of blots of electrophoresis gels of serum showed binding to serum proteins of 200-kDa and four between 75-kDa and 25-kDa for RCA-I and proteins between 150-kDa and 37-kDa for ricin which bound much less strongly. Both ricin and RCA-I bound to purified human immunoglobulin G heavy chain and transferrin but not to α2 HS glycoprotein which contains high amounts of sialic acid. Sambucus nigra agglutinin which binds sialic acid bound to α2 HS glycoprotein and transferrin but not to immunoglobulin G. Ricin bound to human lung A549 proteins of 150 kDa, 100 kDa and several proteins in human cell free salivary fluid between greater than 200 kDa to 25 kDa and binding to these proteins was diminished in the presence of serum.

In fluorescence microscopy, RCA-I bound more strongly than ricin to A549 cells and this binding was diminished in the presence of human serum. RCA-I also bound much more strongly than ricin to human serum proteins coated on microtiter plate wells.

## 1.1 Introduction

Ricin is a highly potent protein toxin and lectin that originates from the plant Ricinus communis, which is readily available throughout the world. Ricin is of great concern and of biomedical and forensic importance for a number of reasons: (a) it has been used by terrorists, criminals and spy agencies inclusive of one incident at a US Senate building and assassination of a Bulgarian dissident in the UK (Schier et al 2007, Audi et al, 2005) (b) there is no cure or antidote (c) ricin poisoning is difficult to diagnose and at high enough doses progresses rapidly to death (Audi J et al. 2005; Poll et al 2009) (d) ricin is banned under both Chemical and Biological Warfare Conventions (Poll et al, 2009). For these reasons, it is important to increase understanding of factors, which influence the ability of ricin to bind and enter cells to cause tissue damage. Ricin kills cells by a process involving binding to cell surface oligosaccharides terminating in galactose followed by internalization and escape from intracellular organelles into the cytoplasm where it enzymatically inactivates ribosomes (Lord et al 1994, Endo et al 1987, Newton et al., 1992). Proteins that are known to bind ricin include IgA (Mantis et al, 2004), the mannose receptor membrane protein (Simmons et al., 1986) the α2 macroglobulin receptor (Cavallero et al, 1995), α2 macroglobulin itself (Ghetie et al., 1991) and the cell surface mannose receptor (Simmons, B.M et al.,1986). One study used affinity chromatography and precipitation in gels to show that Ricinus agglutinin (RCA-I) bound to human IgM, IgA and a fraction of IgG. In that study, RCA-I did not bind to transferrin and α2HS glycoprotein and several other plasma proteins (Harboe et al 1975). Another study identified 28 proteins in intestinal epithelial cell brush border that bound to ricin by using capillary electrophoresis and mass spectrometry. Seven of these were cell surface integral membrane proteins but many were nuclear and cytoplasmic proteins that do not contain galactosylated oligosaccharides that bind to ricin (Liu et al, 2009). Interaction of ricin with proteins in blood is one factor that could affect the binding of ricin to cells. Several plasma glycoproteins exist in mg/ml concentrations in blood where they would be in molar excess of poisonous doses of ricin which is able to kill at 50 microgram doses. If plasma glycoproteins contained the galactose-terminated oligosaccharides that bind to ricin, they could affect ricin’s interactions with cells by reducing the amount available that could bind to and enter cells. However, knowledge can be increased of the specific molecular identities of ricin-binding proteins in human blood. The relatively non-toxic lectin RCA-I which is also found in Ricinus communis is reported to have polypeptide sequence homology and similar oligosaccharide binding to those of ricin (Audi et al, 2005, Lord et al, 1994). Study of RCA-I could help clarify ricin interaction with proteins in blood and other targets by using a less toxic lectin.

### 1.2 Hypothesis

It is here hypothesized that certain serum proteins such as transferrin and immunnoglobulin G would bind to ricin and RCA-I as certain of these proteins contain nonsialylated terminal galactose oligosaccharides (Clerc et al, 2012) to varying extents. It was also hypothesized that certain lung A549 and adrenal SW-13 cell lines and saliva cell free proteins would bind to ricin and RCA-I.

### 1.3 Objectives

Apart from ricin, the plant Ricinus communis contains a lectin RCA-I that is reported to have similar or identical carbohydrate binding specificity to ricin. The binding of RCA-I to cells and serum will also be studied since it might be used as a less toxic surrogate for clarifying which proteins ricin might interact with.

The objectives of the present study were to (1) characterize the interactions of ricin with human serum proteins and identify proteins that bound to ricin in serum, A549 and SW-13 cells and human cell free salivary fluid (2) assess the degree to which proteins in blood could interfere with ricin binding to target proteins. (3) Compare serum protein binding by ricin to that by RCA-I. (4) Assess the ability of serum to compete binding of RCA-I to cultured cells.

## 2. Materials & Methods

### 2.1 Materials

Biotinylated ricin (biotin-ricin) was obtained from EY Laboratories. Biotinylated RCA-I (biotin-RCA-I), biotinylated Sambucus nigra lectin (biotin-SNA), alkaline phosphatase-conjugated avidin, FITC-conjugated RCA-I and Alexa 488-streptavidin were obtained from Vector Laboratories. Human transferrin was purchased from Calbiochem. Human IgG was obtained from Sigma. Human α_2_ HSGP was purchased from the Binding Site. Human serum was obtained from Zen-Bio. Alkaline phosphatase microwell substrate was provided by Kirkegaard and Perry Laboratories. A549 and SW-13 cells and their culture media were obtained from ATCC. Flat bottomed polystyrene microtiter plates (part no. 655101) were obtained from Greiner Bio-one. Reagents and buffers for SDS-PAGE, Kaleidoscope Precision Plus pre-stained molecular mass markers, Tween-20, non-fat milk powder, mini trans-blot electrophoretic transfer cells and nitrocellulose were obtained from Biorad Laboratories. Phosphate buffered saline tablets were purchased from Amresco. Thermo Scientific Nunc LabTek Chamber eight well slides for fluorescence staining of cultured adherent cells were obtained from Daigger Scientific. The Kaleidagraph graphing and data analysis software was used to create graphs and derive basic descriptive statistical analysis of data.

### 2.2 Methods

#### 2.2.1 Microtiter plate lectin binding assays

To do the microtiter plate assay for the binding of different concentrations of biotinylated RCA-I and ricin to a fixed concentration of serum on microtiter plates, human serum was initially diluted 512-fold in the buffer 2 mM Na_2_CO_3_, 8 mM NaHCO_3_, 150 mM NaCl, pH 10.5. This solution was added as 100 µL aliquots in duplicate into wells of a 96 well microtiter plate. The first two columns of the plate were blank wells to which were added buffer only. Incubation was done for 16h at 4 degrees centigrade to allow serum proteins to stick to the microtiter plate wells. The wells were then emptied and rinsed twice with 300 µL per well of phosphate buffered saline, pH 7.4. To block free binding sites on the microtiter plate, 300 µL of 2.5% nonfat milk in PBS was added to all wells and incubated for 2h at 25 degrees centigrade. The wells were then emptied and washed four times with PBS, 0.1% Tween-20.

For lectin binding, the microtiter wells were incubated for 60 min at 25° C with 100 µl of varying concentrations of ricin or RCA-I, 1 µg/ml to 8ng/ml that were produced by 2 fold serial dilutions in PBS 0.1 % Tween 20. The wells were emptied and washed 5 times with 300 µl of PBS, 0.1% Tween 20. The wells were then incubated with 100 µl of 1µg/ml alkaline phosphatase conjugated Streptavidin for 30 min at 25 ^0^C. The microtiter plate wells were emptied and washed 5 times with PBS Tween 20. The plates were incubated for 15 minutes with 100 µL per well of alkaline phosphatase microwell substrate from Kirkegaard and Perry Laboratories. The absorbencies were measured at 490 nm by using a Biorad microtiter plate reader model 480.

The binding of 1µg/ml biotinylated RCA-I and ricin to 24 different samples of human serum was measured by diluting sera 1024-fold in 2 mM Na_2_CO_3_, 8 mM NaHCO_3_, 150 mM NaCl, pH 10.5. The coating of microtiter plate wells, incubations, blocking and detection and measurement of binding by streptavidin alkaline phosphatase and substrate were done as described above.

#### 2.2.2 Lectin blot: lectin staining of electrotransfers of SDS PAGE (sodium dodecyl sulfate polyacrylamide electrophoresis) gels

Human sera diluted twenty-fold in PBS and purified human transferrin, immunogobulin G and α2 HSGP (1 mg/ml in PBS) were prepared for SDS PAGE by adding an equal volume of SDS gel sample buffer containing 10mM dithiothreitol (reducing agent) and boiling for 2 min. For sample preparation of A549, SW-13 and buccal cells for SDS –PAGE, the cells at a density of 5 × 10^6^ cells/mL were mixed with an equal volume of SDS sample buffer and reducing agent and heated. The samples were centrifuged for 10 minutes in a microcentrifuge at 10,000 g and the supernatants used for electrophoresis. Twenty-five microliters of the SDS solubilized proteins were subjected to SDS PAGE by the method of Laemmli, 1970 by using precast 4-20% gradient gels from Biorad. In order to perform lectin blotting after SDS PAGE, proteins were electro-transferred to nitrocellulose as previously described (Vanderpuye et al 1991). Nitrocellulose transfers (blots) of the SDS PAGE gels were blocked for 2hrs with 2.5% nonfat milk from Biorad. Transfers were then incubated for 1 hr with 1µg/ml of biotin-conjugated lectin: one of ricin, RCA-I, or Sambucus nigra agglutinin (SNA) in PBS, 0.1% Tween 20. The transfers were then washed 5 times with PBS, 0.1% Tween 20 and incubated 30 min in 1µg/ml alkaline-phosphatase-conjugated streptavidin in the same buffer. After washing 5 times with PBS, 0.1% Tween 20, the transfers were developed for 10-20 minutes with alkaline phosphatase membrane substrate 5-bromo-4 chloro-3 indoyl phosphate and nitro blue tetrazolium from Kirkegaard and Perry Laboratories. The lectin blotting used here is similar in principle and procedure to that described by Cao et al. 2013.

#### 2.2.3 Cell culture and Fluorescence Microscopy

The SW-13 adrenal cancer cell line was cultured by following the American Type Culture Collections’ instructions. Cells were cultured in Leibovitz’s L-15 medium containing 10% fetal bovine serum at 37°C in a 5% CO_2_ atmosphere.

The A549 human lung cancer cell line was cultured by following the American Type Culture Collections’ instructions. Cells were cultured in F-12K medium containing 10% fetal bovine serum at 37°C in a 5% CO_2_ atmosphere. Staining of adherent A549 cells with FITC-conjugated RCA-I was performed with cells grown on microscope slides in the Chamber-Slide system.

The cells in the chamber slide wells were washed four times with 500 µL PBS for each well. For incubations in the absence of serum, 200 µL PBS containing 1µg/mL RCA-I and 100µg/ml Hoechst 33342 were added to duplicate chamber slide wells. For incubations in the presence of serum, 100 µL of human serum was mixed with 100 µL PBS and a final concentration of 1µg/mL RCA-I and 100µg/ml Hoechst 33342 and added to duplicate chamber slide wells.

Incubation was performed for 30 min at 25° Centigrade. The chamber slide wells were then washed five times with 500 µL PBS. The wells were removed, and coverslips added over the cells which were viewed by using a fluorescence microscope under the FITC and DAPI filters and phase contrast conditions.

Staining of adherent A549 cells with ricin was done by the same protocol used for RCA-I except that incubation with biotin-conjugated ricin was used and the cells were then washed and incubated with avidin conjugated to the green-fluorescent dye Alexa to detect biotinylated ricin. Incubations were likewise done for 30 minutes and four washes with 500 µL of PBS done after each incubation.

Fluorescence microscopy was done with an Olympus BX-41 microscope with an attached Olympus Q color 5 camera (UCMAD3) and pictures were captured by using Image J (Image Pro) software. Objectives were Olympus Plan 40X.0.65 Ph2, ∞/0.17 and Olympus Plan 10X/0.25 ∞/-.

## 3. Results

### 3.1 Serum proteins bound by ricin, Ricinus communis I (RCA-I) and Sambucus nigra agglutinin (SNA) after SDS gel electrophoresis and electroblotting (Western blotting)

The proteins stained by Coommassie blue for serum samples from six different individuals and for α2 HSGP, transferrin and Immunoglobulin G are shown in Fig 1A. The serum protein samples are essentially identical. The positions of transferrin (a) and Immunoglobulin G heavy chain (b) and a protein close to 25 kDa are indicated. The α2 HSGP migrates close to the position of immunoglobulin G and stains very poorly with Coommassie Blue dye owing to its very high level of glycosylation (lane 7, Fig. 1A).

**Fig. 1A.**
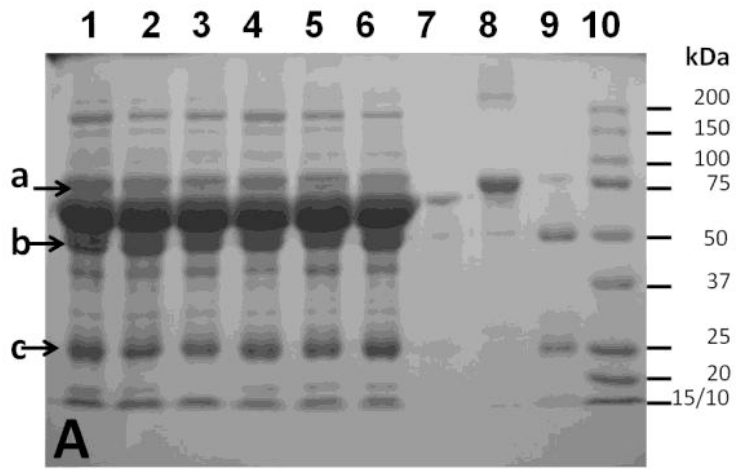
Coomassie stained SDS polyacrylamide gel of serum samples, alpha 2-HS glycoprotein, transferrin and Immunoglobulin G.

**Fig. 1 B.**
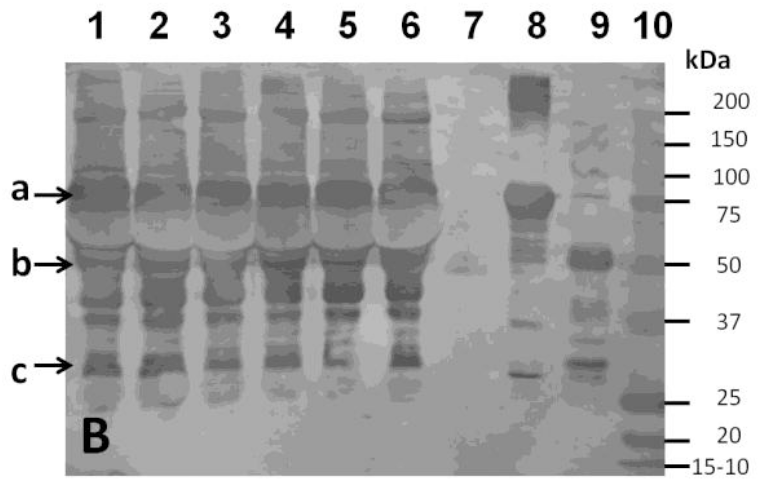
RCA-I lectin blot of SDS gel of various serum samples.

**Fig. 1 C.**
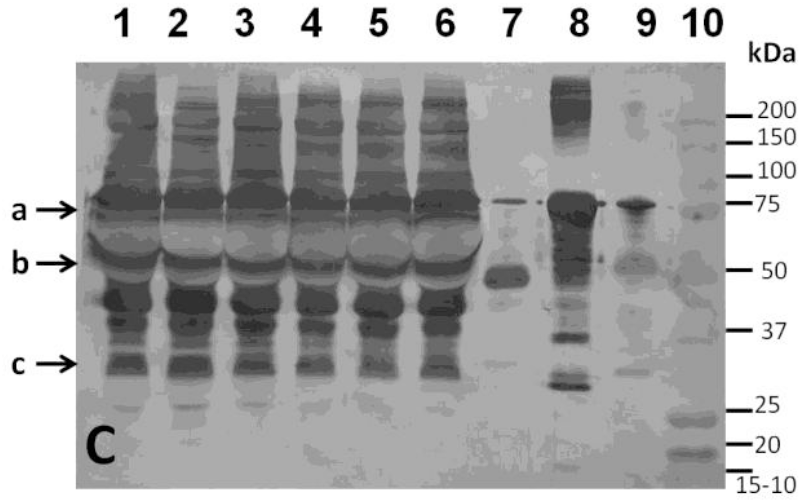
Sambucus nigra lectin blot of SDS gel of various serum samples.

**Fig. 1 D.**
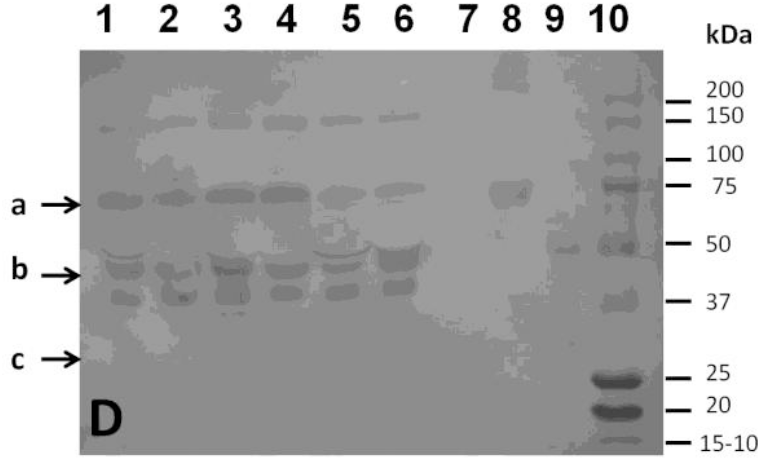
Ricin lectin blot of SDS gel of various serum samples.

The same protein samples were stained on nitrocellulose transfers by using RCA-I. This lectin produced strong staining and the same pattern for all six serum samples. The main serum proteins stained by RCA-I (Figure 1.B.) had relative molecular masses of 200 kDa, 100kDa, about 80 kDa-70 kDa and four bands from 50 kDa to 30 kDa (c indicates the ∼ 30 kDa protein). RCA-I bound to transferrin and immunoglobulin G heavy chain but did not bind to α2 HSGP. The lectin SNA produced very strong staining of all serum samples that resembled that produced by RCA-I (Figure 1.C.). Two serum proteins between 200 kDA and 100 kDA were more strongly stained by SNA. SNA stained α2 HSGP unlike RCA-I and also stained transferrin and contaminating high and low molecular weight material in this sample. SNA only very weakly stained 50kDa material in the immunoglobulin G preparation.

Ricin staining of all six serum protein samples was much weaker than that of RCA-I and SNA (figure 1.D.). Ricin bound to four serum proteins including a serum protein close to but less than 200 kDa and proteins at the positions of transferrin and immunoglobulin G heavy chain and protein just below the immunoglobulin G heavy chain. Ricin did not bind to α2 HSGP but did bind to transferrin and immunoglobulin G heavy chain (lanes 8 and 9, respectively, Fig. 1 D.

### 3.2 Ricin staining of blots of SDS PAGE gels of proteins from A549 cells, SW-13 cells, buccal cells and serum in the presence and absence of serum

Ricin bound to A549 cell proteins with reduced disulfide bonds in the region of 120 kDa, 100 kDa, 75 kDA and 20 kDa. In SW-13 cells, ricin bound less strongly overall and recognized a protein less than 20 kDa. Salivary fluid cell free proteins were strongly stained by ricin including ones of greater than 200 kDa, 120 kDa, and the region from 75 kDa to 20 kDa. For the serum sample used, proteins of 120 kDa, 75 kDa and 50 kDa were bound by ricin (Figure 2, left panel).

**Fig. 2.**
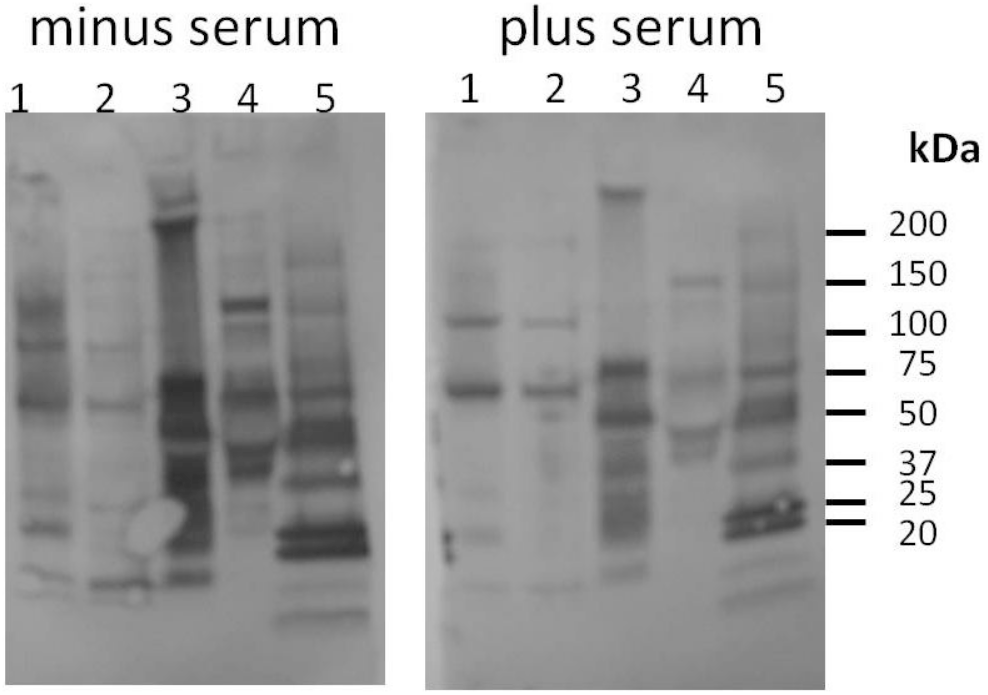
Ricin blot of A549 cells, SW-13 cells, human salivary fluid, human serum in the absence and presence of serum. Lanes, 1-A549 cells, 2- SW-13 cells, 3-human salivary fluid (minus buccal cells), 4- human serum, 5-molecular mass standards

The staining of proteins from A549 cells, SW-13 cells, buccal cells and serum by ricin was decreased when serum was co-incubated with the ricin during staining of the blots (Figure 2 right panel). This was less evident however for the less strongly stained SW-13 cell proteins.

### 3.3. RCA-I binds better to serum proteins than does ricin in a microtiter plate assay

Binding to a fixed concentration of serum proteins was compared between RCA-I and ricin at concentrations from 8ng/ml to 1µg/ml. (Figure 3). Binding of RCA-I to serum increased over the range 8ng/ml to 0.2 µg/ml of RCA-I and then decreased. This is similar to the prozone or hook effect for antibody binding to antigen in microtiter plate assays (Butch, A. 2000). Binding of ricin to serum very gradually increased over the range 8ng/ml to 0.2mg/ml and then reached a plateau. The absorbance produced by the binding of RCA-I to serum was much higher than that of ricin at all concentrations of these lectins. The maximum absorbance for RCA-I binding to serum was 0.7 and that of ricin was 0.04 and thus there was 17.5-fold greater binding of RCA-I than ricin to serum when both were at a concentration of 0.2 mg/ml.

**Fig. 3.**
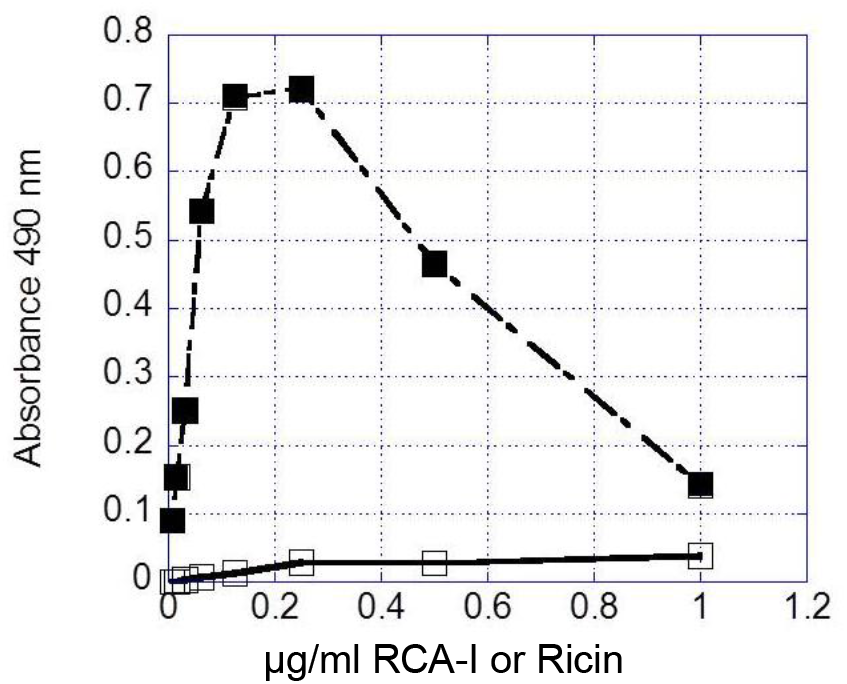
RCA-I binds better than ricin in a microtiter plate assay titration of binding to serum proteins. Solid squares-RCA-1 binding; open squares – ricin binding µg/ml RCA-I or Ricin

The binding of ricin and RCA-I was also compared for the same set of 24 different serum samples (Fig. 4.). (These serum samples were 2-fold greater diluted than the one used for Figure 3.) For all serum samples, the binding of RCA-I was much greater than that of ricin. The average and the median absorbencies for RCA-I binding were0.657 and 0.648 respectively (S.D – 0.065 and SEM - 0.013). The binding of ricin was reflected by an average absorbance of 0.033 and median of 0.34 (SD. – 0.022 and SEM 0.004). The average absorbance for RCA-I binding to the 24 different serum samples was therefore 19.9-fold higher than that of ricin.

**Fig. 4.**
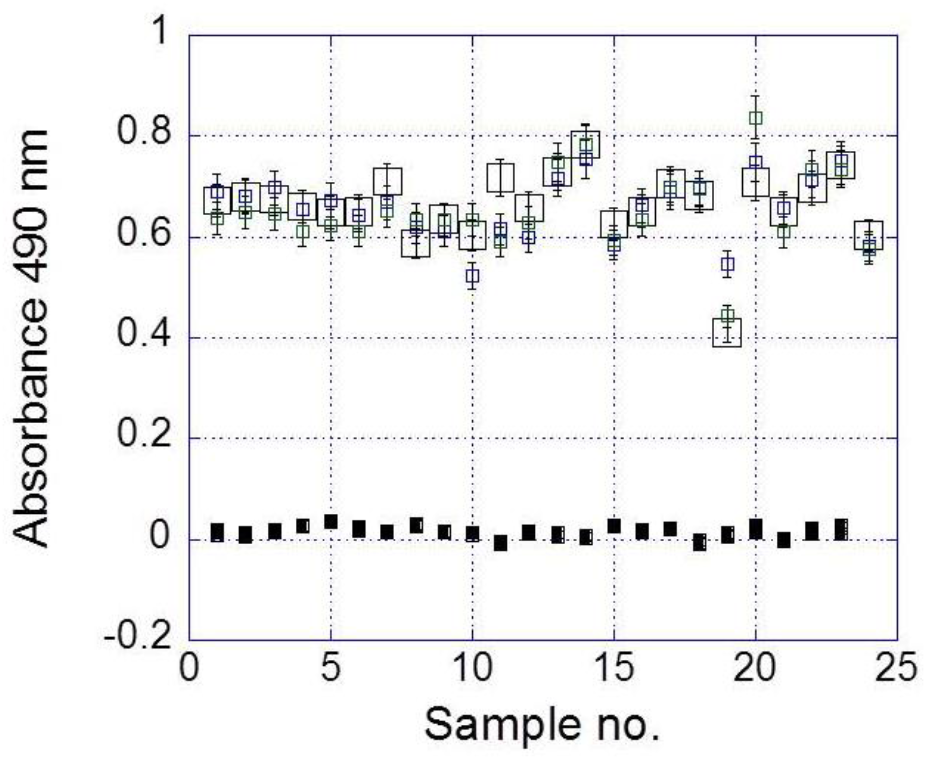
Scatter plot of microtiter plate binding of ricin and RCA-I to 25 different serum samples. Solid squares-RCA-1 binding; open squares – ricin binding

### 3.4. RCA-I binds to human A549 lung cells in fluorescence microscopy and its binding is inhibited by human serum

Biotinylated RCA-I and Hoechst 33342 nuclear dye were used to stain adherent A549 cells, and the results visualized by fluorescence microscopy and phase contrast (Figure 5 A-C). The biotinylated RCA-I bound to the surface of A549 cells and also produced diffuse and punctate cytoplasmic staining. RCA-I also bound quite distinctively to areas of cell-cell contact as bands (Figure 5A). The nuclei of the cells in the same slide as Figure 5A and the outlines of the same cells are shown by Hoechst 3342 nuclear dye staining (Figure 5B) and phase contrast imaging (Figure 5C). Comparing the images from Figure 5A to C shows that most of the A549 cells bound RCA-I.

**Fig 5A.**
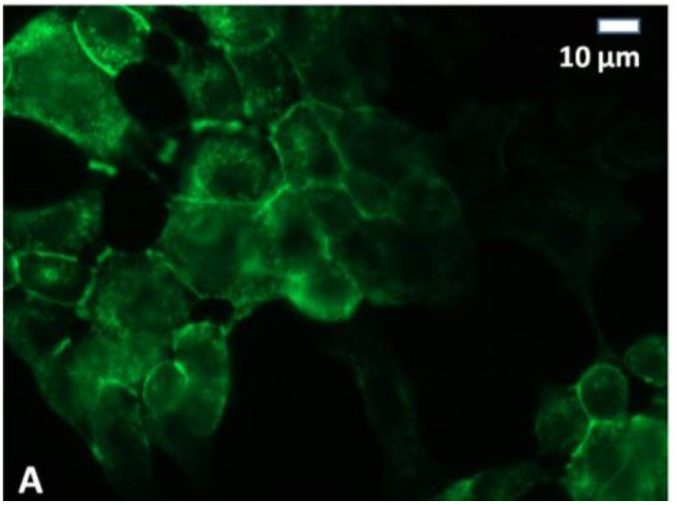
RCA-I staining fluorescence, binding to A549 cells minus serum.

**Fig 5 B.**
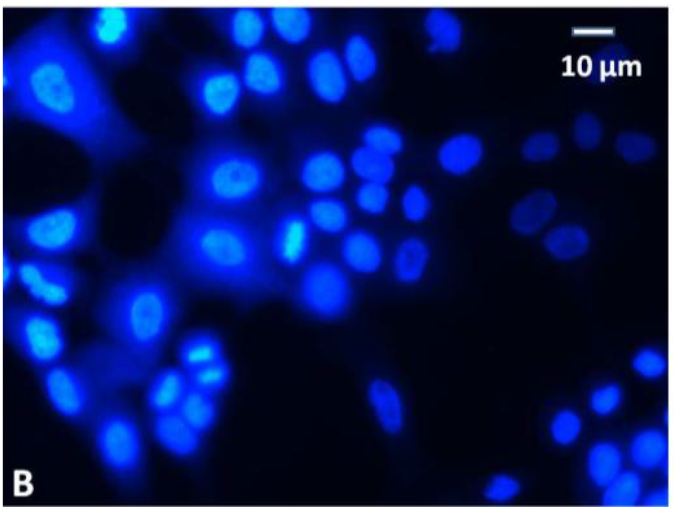
Hoechst nuclear fluorescence, A549 cells incubated with RCA-I minus serum.

**Fig 5 C.**
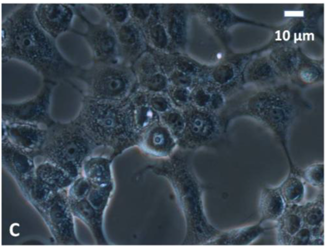
phase contrast microscopy, A549 cells incubated with RCA-I minus serum.

**Fig 5 D.**
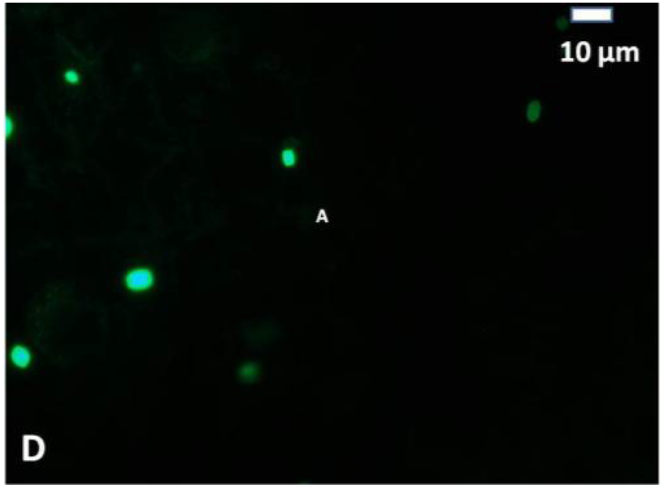
RCA-I staining, fluorescence, binding to A549 cells plus serum.

**Fig 5 E.**
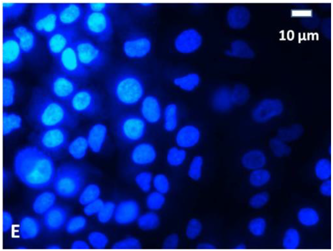
Hoechst 33342 nuclear fluorescence, A549 cells incubated with RCA-I plus serum.

**Fig 5 F.**
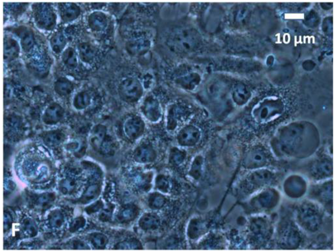
phase contrast microscopy, A549 cells incubated with RCA-I plus plus serum.

Fluorescence and phase imaging are shown in Figures 5.D-F. for a slide of the same A549 cells incubated with RCA-I and Hoechst 3342 in the presence of 50% human serum. The staining of A549 cells by RCA-I was abolished or greatly decreased by co-incubation of serum with RCA-I (Figure 5D) The Hoechst nuclear staining (Figure 5E) and phase contrast imaging (Figure 5F) showed that cells with nuclei and cell borders and cytoplasmic structures were present. Similar results were obtained with FITC-labeled RCA-I.

### 3.5. Fluorescence microscopy of human A549 lung cells incubated with ricin in the presence and absence of human serum

Since the binding of ricin to proteins from A549 cells was detected by Western blotting, the binding of ricin to A549 cells cultured on microscope slides was examined by fluorescence microscopy (Figs 6A-F). The same cells incubated with biotinylated ricin and Hoechst 33342 in the absence of serum are shown in Fig. 6A with filters for green fluorescence to detect binding of Alexa conjugated to avidin to biotinylated ricin. In Fig. 6B, imaging of these cells was done with blue filters to detect Hoechst staining of nuclei. Phase contrast imaging of the same cells is shown in Fig 6C.

**Fig 6.A.**
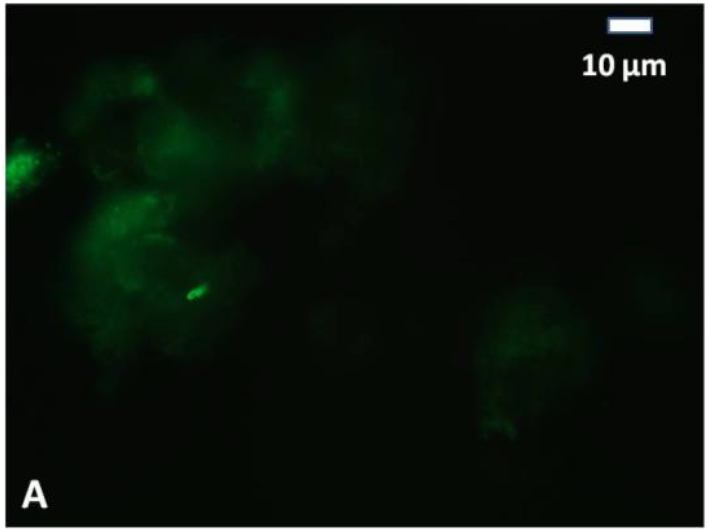
FITC Fluorescence microscopy, ricin incubation with A549 cells minus serum.

**Fig 6 B.**
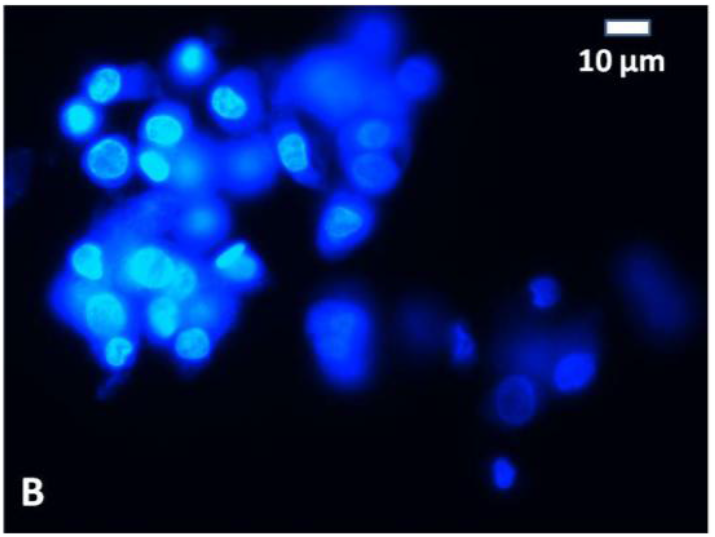
Hoechst nuclear fluorescence microscopy, ricin incubation with A549 cells minus serum.

**Fig 6 C.**
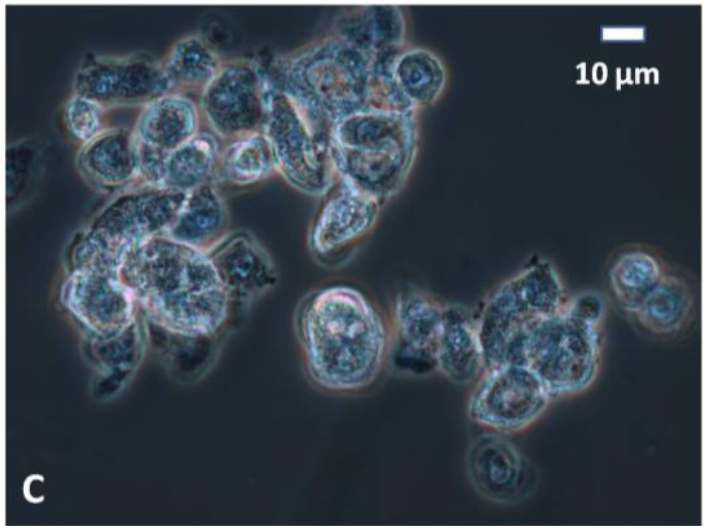
Phase contrast microscopy ricin incubation with A549 cells minus serum.

**Fig 6 D.**
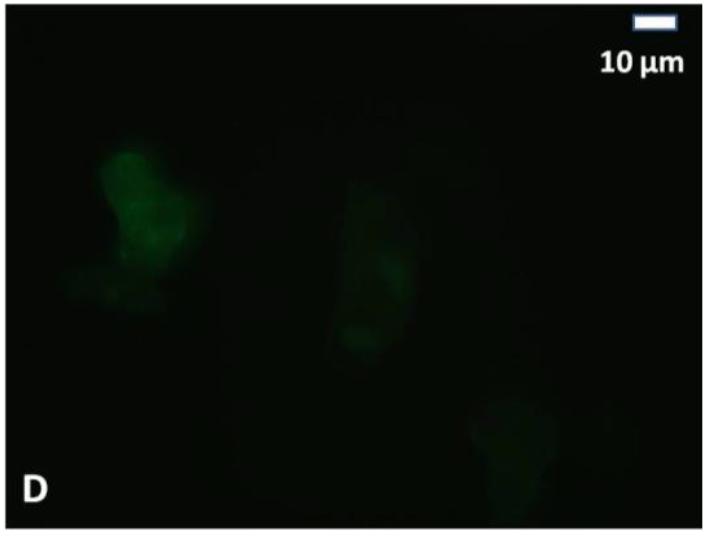
FITC Fluorescence microscopy, ricin incubation with A549 cells plus serum.

**Fig 6 E.**
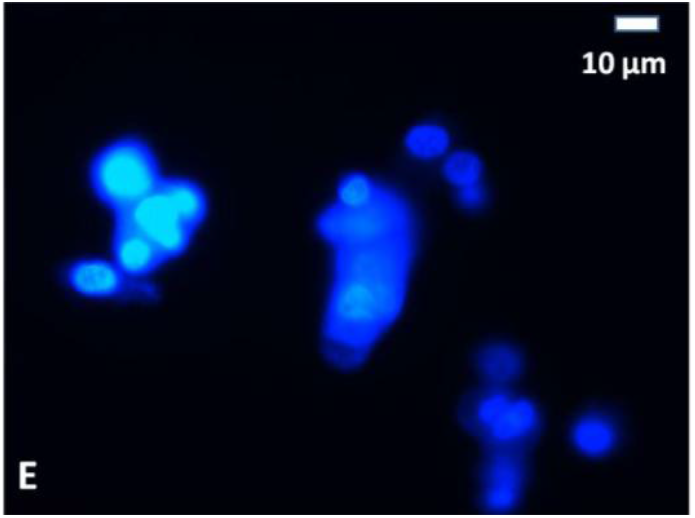
Hoechst nuclear fluorescence microscopy, ricin incubation with A549 cells plus serum.

**Fig 6 F.**
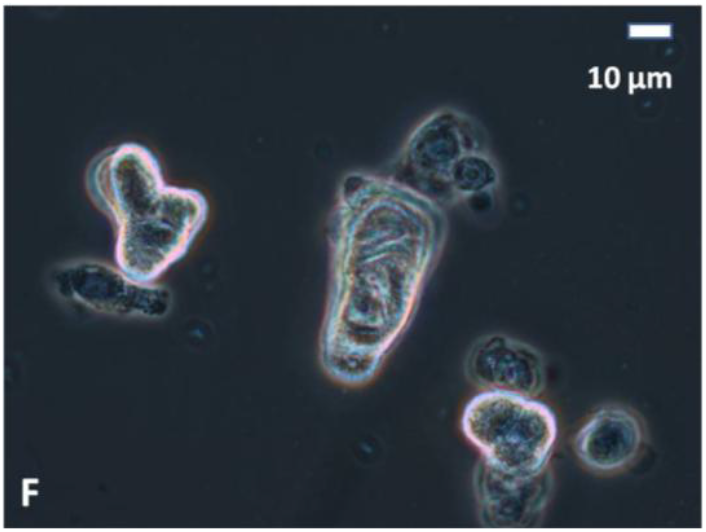
Phase contrast microscopy, ricin incubation with A549 cells plus serum.

The results showed only weak binding of ricin to A549 cells in Fig 6A although Figs 6B and 6C showed the presence of nuclei and cells. Similar results were obtained for ricin incubation of A549 cells in the presence of serum. The binding of ricin to A549 cells was much very much weaker and a different pattern to that observed with RCA-1 at the same concentration.

## 4. Discussion

This study examined the binding of ricin and RCA-I to serum glycoproteins, human lung A549 cells and SW-13 cells and salivary cell free fluid. The hypothesis that human serum glycoproteins can inhibit ricin and RCA-I binding was tested. RCA-I binding to proteins was also examined since it is highly related to ricin in structure and has been reported to have the same sugar specificity (Roberts et al, 1985; Debray et al, 1981).

This study made four main findings. First, ricin and RCA-I bind purified human transferrin and the 50,000 kDa heavy chain of immunoglobulin G but not α2HS -glycoprotein in electrophoretic gel blotting analyses. A major protein in serum bound by both ricin and RCA-I had a molecular mass of approximately 200 kDa under reducing conditions. Ricin bound weakly to purified immunoglobulin G and transferrin compared to RCA-I and to proteins in serum of 200 kDa, 75 kDa,50 kDa and between 50kDa and 37 kDa. It did not bind to α2 HS-glycoprotein as expected because oligosaccharides on this protein lack terminal galactose and mainly have terminal sialic acid (Wilson et al., 2002, Clerc et. al. 2012). RCA-I bound strongly to proteins in serum of 200 kDa, 100 kDa, 80-70 kDa and 50-30 kDa. Transferrin and the heavy chain of IgG have both been characterized as containing terminal beta-linked galactose (Bondt et al., 2014, Vidarsson et al., 2014, Clerc et. al. 2012). RCA-I did not bind to α2 HS-glycoprotein which is reported to contain large amounts of sialic acid as a terminal sugar (Vidarsson et. al., 2014, Clerc et. al. 2012). In line with these results, consistent with its reported specificity for terminal sialic acid, the lectin Sambucus nigra agglutinin bound very strongly to α2 HS-glycoprotein and transferrin but not to immunoglobulin G. Immunoglobulin G, transferrin and α2 HS-glycoprotein are present in large or significant amounts (mg/ml) in blood (serum and plasma) so that potentially if they bound to ricin they would be expected to diminish its toxicity.

The results did show that despite ricin being structurally related to RCA-I and its reported specificity for oligosaccharides that have terminal galactose, ricin and RCA-I have some differences in their binding to serum proteins.

Reports are lacking on the direct observation of which serum proteins are bound by ricin. There have been more studies on the binding of proteins in blood (plasma/serum) to RCA-I than on the binding of proteins to ricin.

The identification of transferrin, immunoglobulin G (IgG) as serum proteins bound by ricin and RCA-I was consistent with the oligosaccharide structures reported to be present on these proteins and the sugar binding specificities of ricin and RCA-I (Bondt et al,. 2014., DeBray et al., 1981, Vidarsson et al, 2014). Oligosaccharides on transferrin include bi-antennary N-linked mono-sialylated structures with terminal galactose on one chain (Wilson et. al., 2002, Clerc, F et al.).

Oligosaccharides on IgG include mono-galactosylated hybrid and bi-antennary N-linked structures with terminal galactose (Vidarsson et al., 2014, Bondt et. al, 2014, Clerc, F et al.). These types of N-linked oligosaccharide structures on transferrin and IgG are ones that are bound by ricin and RCA-I.

The 200 kDa serum protein that was bound by ricin and RCA-I could possibly have been alpha 2-macroglobulin which is highly glycosylated and has bi-antennary mono-sialylated N-linked structures (Lin et al., 2012, Clerc, F et al.). Alpha 2-macroglobulin was also reported to bind to ricin by mechanisms other than lectin reactivity with oligosaccharides (Ghetie et al., 1991). The specificity of the reactivity of serum proteins with ricin and RCA-I by blotting was supported by comparison with their reactivity with SNA. The lectin SNA recognizes oligosaccharides that have terminal sialic acid in α 2-6 linkage to galactose (Shibuya et. al., 1989) while ricin and RCA-I do not bind such structures or do so very weakly but do bind these structures without the sialic acid. SNA did not bind to immunoglobulin G but did bind to transferrin which contains N-linked monosialylated and disialylated bi-antennary structures. The binding of SNA but not ricin and RCA-I to alpha 2 HS-glycoprotein is consistent with the large amounts of sialic acid on this protein, (Wilson et. al., 2002, Clerc, F et al.).

A second finding was that binding of ricin to serum proteins as assessed by ELISA and blotting techniques was much weaker than the binding by RCA-I.

A reason for the difference in binding between RCA-I and ricin may relate to the valency of the serum glycoproteins in terms of the oligosaccharide ligands that they bear and how the spacing between multiple oligosaccharides on the glycoprotein ligand matches the spacing among the binding sites on RCA-I as compared to ricin.

The stronger binding of RCA-I to serum glycoproteins compared to ricin may be also due to the fact that RCA-I is a tetramer whereas ricin is a dimer (Roberts L.M., 1985; Lord et al, 1994).

RCA-I contains two A chains and two B chains in which the B chains each contain two well characterized oligosaccharide-binding sites. Ricin contains one A chain and one B chain which has two main oligosaccharide-binding sites (Lord et al., 1994). The structurally homologous ricin and RCA-I B-chains have extended binding sites that bind oligosaccharides better than monosaccharides. In addition, multiple oligosaccharides at certain distances apart on a glycoprotein are reported to bind the strongest to ricin and RCA-I (Debray et al., 1981). Because of its higher multivalency, RCA-I may bind more strongly than ricin to glycoproteins that have few oligosaccharide chains and to ones in which there is greater spacing between oligosaccharides.

It can thus be envisioned that for a multivalent ligand, RCA-I would be more likely than ricin to remain bound because of the larger number of binding sites on RCA-I and the increased chance that one of the sites would encounter a glycoprotein oligosaccharide ligand.

A third finding was that ricin bound to different proteins separated by SDS gel electrophoresis in lectin blots of A549 cells, SW-13 cells, human salivary cell-free fluid and as a control serum and the binding could be decreased by human serum. There were no direct matches in the molecular weights of the ricin-binding proteins in the different samples except that 150 kDa proteins in serum and A549 cells both bound ricin. The results suggest that glycoproteins in blood could bind ricin and affect its binding to cells.

The fourth finding was that serum inhibited RCA-I binding to A549 cells when assessed by fluorescence microscopy. RCA-I was very efficiently inhibited from binding to A549 cells by serum in immunofluorescence microscopy experiments indicating that serum can prevent binding of lectins to cells and suggesting that this could happen in vivo. Since ricin binds to serum proteins, it is possible that in cases of ricin poisoning complexes of ricin with IgG and transferrin could occur in blood although this would appear not to eliminate ricin toxicity in vivo. Transferrin is endocytosed via the cell surface transferrin receptor to help deliver iron to cells (Guo, Q et al 2025). Future studies could examine if ricin that is bound to transferrin can enter cells via endocytosis mediated by the transferrin receptor. The alpha 2 macroglobulin receptor binds to the non-lectin chain of ricin and thereby mediates toxicity. Since the lectin component of ricin may bind alpha 2 macroglobulin it could also be studied if alpha 2 macroglobulin complexes of ricin are internalized into cells via the alpha 2 macroglobulin receptor.

Ricin binding to A549 cells was low compared to that of RCA-I at the same concentration by weight even though ricin binding was detected by electroblotting of A549 proteins in polyacrylamide gels. A study in the literature after we completed some of our studies (Smith et al, 2012) described binding of ricin labeled by a fluorescent hydroxy succinimide compound to A549 cells (Jenner et al, 2018).

In summary, this study showed that transferrin, immunoglobulin G and a close to 200 kDa molecular weight protein under reducing conditions are among the serum proteins that bind to ricin. Proteins that bound to ricin in human lung A549 cells and salivary fluid were also detected. Serum proteins can decrease ricin binding to cells as assessed by lectin blot and immunofluorescence assays. Despite structural homology and similar reported oligosaccharide specificity, RCA-I binds much more strongly to serum proteins than the preparation of ricin used.

## Abbreviations

α_2_2HSGP: α_2_2HS glycoprotein
IgG: immunoglobulin GRCA-I
I: Ricinus communis agglutinin

## Acknowledgements

We acknowledge the Albany State University Center for Undergraduate Research and Title III program for student and faculty undergraduate research awards and the American Society for Cell Biology for a Linkage Fellow Award for cell culture materials. Brandon Walker and Scott Pierce are thanked for help with maintenance of the cell culture and fluorescence microscope facilities.

---

**Fig 1A-D SDS polyacrylamide gel electrophoresis and electroblotting with RCA-1, Sambucus nigra and ricin lectins**

Samples loaded: 1-serum 1, 2-serum 2, 3-serum 3, 4-serum 4, **5-se**rum 5, 6-serum 6, 7-alpha 2-HS glycoprotein, 8-transferrin, 9-Immunoglobulin G, 10 molecular mass standards

